# Comprehensive Mosquito Wing Image Repository for Advancing Research on Geometric Morphometric- and AI-Based Identification

**DOI:** 10.1101/2024.11.13.623340

**Authors:** Kristopher Nolte, Eric Agboli, Gabriela Azambuja Garcia, Athanase Badolo, Norbert Becker, Do Huy Loc, Tarja Viviane Dworrak, Jacqueline Eguchi, Albert Eisenbarth, Rafael Maciel de Freitas, Ange Gatien Doumna-Ndalembouly, Anna Heitmann, Stephanie Jansen, Artur Jöst, Hanna Jöst, Ellen Kiel, Alexandra Meyer, Wolf-Peter Pfitzner, Joy Saathoff, Jonas Schmidt-Chanasit, Tatiana Sulesco, Artin Tokatlian, Thirumalaisamy P. Velavan, Carmen Villacañas de Castro, Magdalena Laura Wehmeyer, Julien Zahouli, Felix Gregor Sauer, Renke Lühken

**Affiliations:** Bernhard Nocht Institute for Tropical Medicine, Hamburg, Germany; Institute for Dipterology (IfD), Speyer, Germany; Center for Organismal Studies (COS), University of Heidelberg, Heidelberg, Germany; Institute of Tropical Medicine, Universitätsklinikum Tübingen, Tübingen, Germany; Vietnamese-German Center for Medical Research, Hanoi, Vietnam; Faculty of Mathematics, Informatics and Natural Sciences, Universität Hamburg, Hamburg; Faculty of Medicine, Duy Tan University, Da Nang, Vietnam; Kommunale Aktionsgemeinschaft zur Bekämpfung der Schnakenplage e. V. (KABS), Speyer, Germany; Laboratório de Mosquitos Transmissores de Hematozoários, Instituto Oswaldo Cruz, Fiocruz, Rio de Janeiro, Brazil; Instituto Nacional de Ciência e Tecnologia em Entomologia Médica, Universidade Federal do Rio de Janeiro, Rio de Janeiro, Brazil; Carl von Ossietzky University, Oldenburg, Germany; Department of Microbiology and Hygiene, Tropical Medical Entomology Unit, Bundeswehr Hospital Hamburg, Hamburg, Germany; Centre de Recherche Médicale de Lambaréné, Lambaréné, Gabon; Laboratory of Fundamental and Applied Entomology, Université Joseph Ki-Zerbo, Ouagadougou, Burkina Faso; Centre Suisse De Recherches Scientifiques En Côte D’Ivoire, Abidjan, Côte d’Ivoire

## Abstract

Accurate identification of mosquito species is essential for effective vector control and mitigation of mosquito-borne disease outbreaks. Traditional morphological identification requires highly specialized personnel and is time-consuming, while molecular techniques can be cost-effective and dependent on comprehensive genetic information. Wing geometric morphometry has emerged as a promising alternative, leveraging detailed geometric measurements of wing shapes and vein patterns to distinguish between species and detect intraspecies variations. This paper presents a curated dataset of 18,104 mosquito wing images, collected from 10,500 mosquito specimens, annotated with extensive meta-information, designed to support research in wing geometric morphometry and the development of machine learning models, ultimately supporting efforts in vector surveillance and research.

## Background & Summary

Mosquitoes are the most important arthropod vectors of pathogens worldwide, e.g. for dengue and Zika virus^1^. Global change, particularly global warming and globalization, facilitates the spread of mosquitoes and their pathogens, increasing the risk of mosquito-borne pathogen transmission in previously unaffected areas^2^. This highlights the need for mosquito monitoring and surveillance methods to develop early prevention measures. Therefore, it is essential to accurately identify mosquito species as the ecology and vector capacity strongly vary among species^3^.

Mosquito species are commonly identified using taxonomic keys based on morphological characters^4^. The morphological species identification can be time-consuming and requires intensive entomological experience, limiting its scalability for large-scale studies^5^. In addition, it has been widely recognized that entomological expertise is declining as traditional taxonomy becomes less central to the biological curriculum^6^. Alternative methods (e.g. DNA barcoding) require specialized equipment and technical expertise^7^, which can be challenging to use in low-resource settings. These limitations highlight the need for complementary approaches and comprehensive datasets to improve species identification accuracy.

Wing geometric morphometrics offers a promising alternative to traditional identification methods of mosquitoes, as it reliably captures interspecific variations^8–10^. This approach utilizes the coordinates of anatomical features of the wing to analyse shape differences between species. Furthermore, the method can be used to detect subtle intraspecific differences, providing valuable insights into population structure, breeding conditions, or fecundity^11–13^. This makes wing geometric morphometrics a versatile and powerful tool for entomological research, offering a scalable and cost-effective solution for large-scale studies in mosquito biology and vector control. However, the analysis is based on the manual setting of selected landmarks on the mosquito wing image, a time-consuming and observer-biased process^14^.

Advances in computer vision, utilizing image processing and machine learning techniques, offer new pathways by automating the species and landmark identification process^15^. Convolutional Neural Networks (CNN) have demonstrated a high potential in accurately identifying mosquito species based on images of either the entire body^16–18^ or solely the wings^19,20^. For instance, Sauer et al. reported that a CNN model trained on wing images achieved a 91% macro-F1 score in classifying seven *Aedes* species^19^. Additionally, segmentation and regression models were used to automatically identify landmarks in Diptera. For example, Geldenhuys et al. (2023) demonstrated this for Tse Tse flies (*Glossina* spp.), training their model on 14,534 pairs of wings of two different species and achieving automated landmark detection with a mean distance error of 3.43 pixels per landmark^21^.

Despite these advancements, the public availability of large high-quality datasets remains a bottleneck for the widespread application of geometric morphometrics and machine learning techniques. Existing wing datasets are often limited in size or scope, hindering the development of robust and generalizable models^22,23^. Herein, we present a dataset with a comprehensive collection of diverse wing images from 72 mosquito taxa from 12 countries on five continents collected between 2008 and 2024. This dataset supports traditional morphometric studies but also enables the application of advanced machine learning techniques, creating opportunities for new insights and innovations in mosquito surveillance and research. By providing this information to the scientific community, we aim to accelerate the progress of research in mosquito biology, vector control, and disease prevention.

## Methods

### Mosquito Sampling

The wing dataset is composed of images consolidated from various experimental and field studies. This study consolidates the efforts of 22 research projects conducted from 2008 to 2024, incorporating a total of 10,500 mosquito specimens from 12 countries across 5 continents (Fig. 1). Various methods were used to collect the mosquitoes, including CO_2_-baited traps (n=4,614), aspirators (1,630), ovitraps (308), egg-raft collection (1,683), and rearing from breeding facilities (2,113). Most of the sampled mosquitoes were identified as female (n=9,049), with a smaller proportion being male (1,425). Species identification was primarily conducted using morphological methods (n=4,658)^4^ or molecular techniques such as COI/nad4 gene barcoding (1,914)^24^, ITS2 gene barcoding for the identification of *Anopheles* species (621)^25^ and qPCR targeting CQ11 and ACE2 genes to identify the taxa morphologically identified as *Culex pipiens* s.l./*torrentium* (2,092)^26^. The remaining samples were identified based on their association with a laboratory colony (1,839).

**Figure 1.**
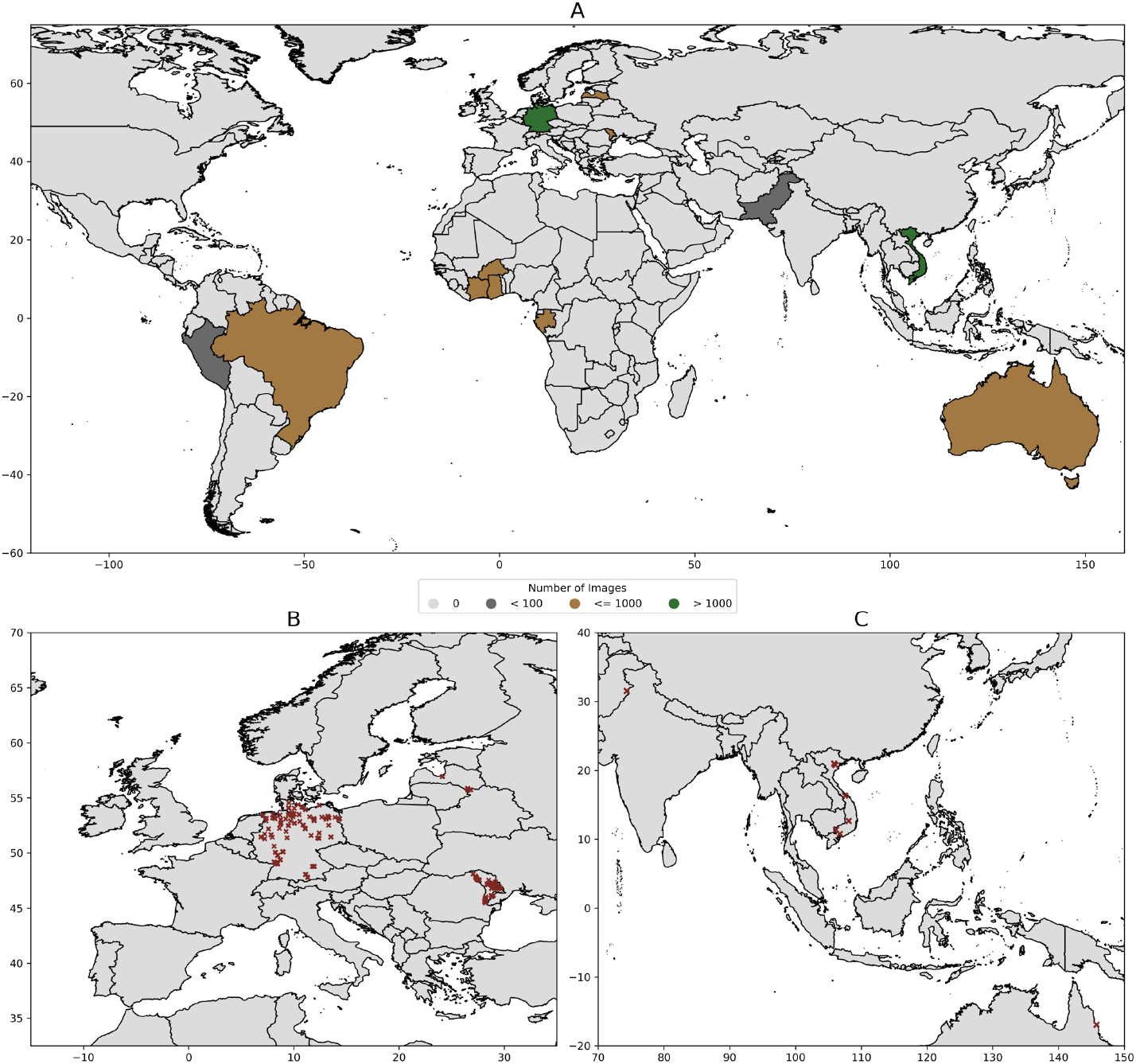
Geographic distribution of images in the dataset. Countries colour-coded by the number of images (1a). Panel 1b shows the mosquito sampling locations in Europe and 1c sampling locations in Southeast Asia. We included both Europe and Asia in the figure due to their higher variance in sampling locations compared to Africa and South America.

The complete wing dataset comprises specimens from nine genera: *Culex* (n=3,980), *Aedes* (5,029), *Anopheles* (1,135), *Coquillettidia* (141), *Culiseta* (158), *Uranotaenia* (1), *Armigeres* (49), *Mansonia* (6), and *Toxorhynchites* (1). Many mosquito species are difficult or impossible to identify based solely on morphology. However, the dataset includes specimens from such groups, which were identified using morphological characteristics alone. Thus, to present the taxonomic information in a machine-readable format, we developed a hierarchical system of taxonomic levels that also describes the uncertainties in species identification. The first level corresponds to the family, the second level to the genus and the fourth level to species. The third taxonomic level encompasses morphologically very similar species pairs (e.g. *Ae communis*/*Ae. punctor*), species groups (e.g. *Ae. annulipes* group), species complexes (*An. maculipennis* s.l.) or combinations of these aggregated taxa (e.g. *Cx. pipiens* s.l./*Cx. torrentium*). The species names (fourth taxonomic level) for these specimens were assigned only when the identification was confirmed through molecular assays. Information on subspecies or biotypes is presented under the fifth taxonomic level.

### Wing Preparation and Image Capture

Wings were removed from mosquitoes with tweezers under a stereo microscope. The wings were placed on a microscopic slide and embedded in Euparal (Carl Roth, Karlsruhe, Germany) with a cover slide for long-term storage. Detailed instructions on the wing removal process are provided in the supplementary material (see supplementary material: wing_removal_instructions.pdf). In total 18,104 images were captured using different stereomicroscopes and a smartphone with an attached macro-lens. Most images (n = 12,462) were captured using the Olympus SZ61 (Olympus, Tokyo, Japan) in conjunction with the Olympus DP23 camera (Olympus, Tokyo, Japan), followed by 3,577 images with the Leica M205c microscope (Leica Microsystems, Wetzlar, Germany) and 1,685 images using an iPhone SE 3rd generation (Apple Inc., Cupertino, USA) in combination with a macro-lens taken at 24x magnification (Apexel-24XMH, Apexel, Shenzhen, China). The images were captured in TIF format, with resolutions of 3024×3024 for smartphone images, 3088×2076 for Olympus DP23 images and 2560×1920 for images captured using the Leica M205c. Imaging settings were not standardized and thus varied between image collection projects in parameters not further recorded, e.g. exposure time or lighting conditions. The images captured with the stereomicroscopes are displayed with a scale providing a reference for wing size measurements such as wing length.

### Data Records

The image dataset was uploaded to the Bioimage Archive and published under a CC-BY 4.0 license (S-BIAD1478, https://doi.org/10.6019/S-BIAD1478). Images are organized in a hierarchical folder structure with separate folders per 2^nd^ taxonomic level, i.e. genus. The images are named according to the following template and comprehensive metadata (e.g., sampling location or image capture device) is provided for each image (Table 2).

**Table 1:** Distribution of specimens and images separated by taxonomic levels.

| 1. Taxonomic Level | 2. Taxonomic Level | 3. Taxonomic Level | 4. Taxonomic Level | 5. Taxonomic Level | Images | Samples |
| --- | --- | --- | --- | --- | --- | --- |
| Family | Genus | Species, Species group, Species complex or Species pairs | Species | Subspecies or biotypes |  |  |
| Culicidae | *Aedes* | - | *aegypti* | - | 2916 | 1300 |
| Culicidae | *Aedes* | - | *albopictus* | - | 1344 | 558 |
| Culicidae | *Aedes* | - | *amesii* | - | 2 | 1 |
| Culicidae | *Aedes* | - | *caspius* | - | 82 | 82 |
| Culicidae | *Aedes* | - | *cataphylla* | - | 148 | 76 |
| Culicidae | *Aedes* | - | *cyprus* | - | 4 | 2 |
| Culicidae | *Aedes* | - | *geniculatus* | - | 24 | 24 |
| Culicidae | *Aedes* | - | *imprimensis* | - | 1 | 1 |
| Culicidae | *Aedes* | - | *intrudens* | - | 17 | 9 |
| Culicidae | *Aedes* | - | *japonicus* | japonicus | 1772 | 689 |
| Culicidae | *Aedes* | - | *koreicus* | - | 1092 | 350 |
| Culicidae | *Aedes* | - | *ostentatio* | - | 19 | 13 |
| Culicidae | *Aedes* | - | *pulchritarsis* | - | 2 | 2 |
| Culicidae | *Aedes* | - | *rossicus* | - | 14 | 14 |
| Culicidae | *Aedes* | - | *rusticus* | - | 229 | 229 |
| Culicidae | *Aedes* | - | *sticticus* | - | 233 | 232 |
| Culicidae | *Aedes* | - | *thailandensis* | - | 2 | 1 |
| Culicidae | *Aedes* | - | *vexans* | - | 455 | 450 |
| Culicidae | *Aedes* | - | *vittatus* | - | 2 | 2 |
| Culicidae | *Aedes* | *annulipes*-group | - | - | 354 | 214 |
| Culicidae | *Aedes* | *cinereus-geminus*-pair | - | - | 339 | 288 |
| Culicidae | *Aedes* | *communis*-punctor-pair | - | - | 580 | 492 |
| Culicidae | *Anopheles* | - | *aconitus* | - | 1 | 1 |
| Culicidae | *Anopheles* | - | *daciae* | - | 18 | 9 |
| Culicidae | *Anopheles* | - | *epiroticus* | - | 6 | 3 |
| Culicidae | *Anopheles* | - | *hyrcanus* | - | 20 | 20 |
| Culicidae | *Anopheles* | - | *messeae* | - | 31 | 16 |
| Culicidae | *Anopheles* | - | *moucheti* | - | 63 | 33 |
| Culicidae | *Anopheles* | - | *paludis* | - | 65 | 33 |
| Culicidae | *Anopheles* | - | *plumbeus* | - | 51 | 47 |
| Culicidae | *Anopheles* | - | *sinensis* | - | 19 | 12 |
| Culicidae | *Anopheles* | - | *stephensi* | - | 251 | 132 |
| Culicidae | *Anopheles* | *claviger-petragnani*-pair | - | - | 136 | 118 |
| Culicidae | *Anopheles* | *coustani*-group | - | - | 80 | 41 |
| Culicidae | *Anopheles* | *gambiae* s.l. | - | - | 42 | 22 |
| Culicidae | *Anopheles* | *maculipennis* s.l. | - | - | 50 | 39 |
| Culicidae | *Anopheles* | *maculipennis* s.l. | *atroparvus* | - | 57 | 57 |
| Culicidae | *Anopheles* | *maculipennis* s.l. | *daciae* | - | 62 | 62 |
| Culicidae | *Anopheles* | *maculipennis* s.l. | *maculipennis* | - | 98 | 98 |
| Culicidae | *Anopheles* | *maculipennis* s.l. | *messeae* | - | 376 | 376 |
| Culicidae | *Anopheles* | *nili* s.l. | - | - | 29 | 15 |
| Culicidae | *Anopheles* | *tessellatus* | - | - | 2 | 1 |
| Culicidae | *Armigeres* | - | *durhami* | - | 1 | 1 |
| Culicidae | *Armigeres* | - | *jugraensis* | - | 1 | 1 |
| Culicidae | *Armigeres* | - | *moultoni* | - | 2 | 1 |
| Culicidae | *Armigeres* | - | *subalbatus* | - | 75 | 46 |
| Culicidae | *Coquillettidia* | - | *buxtoni* | - | 2 | 2 |
| Culicidae | *Coquillettidia* | - | *crassipes* | - | 63 | 34 |
| Culicidae | *Coquillettidia* | - | *richiardii* | - | 104 | 104 |
| Culicidae | *Coquillettidia* | - | *subalbatus* | - | 2 | 1 |
| Culicidae | *Culex* | - | *bitaeniorhynchus* | - | 8 | 4 |
| Culicidae | *Culex* | - | *brevipalpis* | - | 8 | 5 |
| Culicidae | *Culex* | - | *fuscocephala* | - | 38 | 23 |
| Culicidae | *Culex* | - | *gelidus* | - | 47 | 27 |
| Culicidae | *Culex* | - | *modestus* | - | 146 | 111 |
| Culicidae | *Culex* | - | *nigropunctatus* | - | 9 | 6 |
| Culicidae | *Culex* | - | *orientalis* | - | 4 | 3 |
| Culicidae | *Culex* | *pipiens*<br>s.l.-<br>*torrentium*<br>m-pair | - | - | 2684 | 1885 |
| Culicidae | *Culex* | *pipiens*<br>s.l.-<br>*torrentium*<br>m-pair | *pipiens* | biotype<br>molestus | 353 | 195 |
| Culicidae | *Culex* | *pipiens*<br>s.l.-<br>*torrentium*<br>m-pair | *pipiens* | biotype<br>pipiens | 1913 | 978 |
| Culicidae | *Culex* | *pipiens*<br>s.l.-<br>*torrentium*<br>m-pair | *quinquefasciatus* | - | 226 | 151 |
| Culicidae | *Culex* | *pipiens*<br>s.l.-<br>*torrentium*<br>m-pair | *torrentium* | - | 462 | 240 |
| Culicidae | *Culex* | *territans-*<br>*hortensis-*<br>pair | - | - | 22 | 22 |
| Culicidae | *Culex* | *vishnui-*<br>group | - | - | 204 | 124 |
| Culicidae | *Culex* | *vishnui-*<br>group | *pseudovishnui* | - | 18 | 9 |
| Culicidae | *Culex* | *vishnui-*<br>group | *tritaeniorhynchus* | - | 385 | 195 |
| Culicidae | *Culex* | *vishnui-*<br>group | *vishnui* | - | 8 | 4 |
| Culicidae | *Culiseta* | - | *alaskaensis* | - | 2 | 1 |
| Culicidae | *Culiseta* | *annulata-*<br>*subochrea-*<br>pair | - | - | 50 | 50 |
| Culicidae | *Culiseta* | *morsitans*<br>-<br>*fumipennis*<br>s-pair | - | - | 168 | 107 |
| Culicidae | *Mansonia* | - | *indiana* | - | 3 | 2 |
| Culicidae | *Mansonia* | - | *uniformis* | - | 6 | 4 |
| Culicidae | *Toxorhynchites* | *splendens* | - | - | 1 | 1 |
| Culicidae | *Uranotaenia* | - | *unguiculata* | - | 1 | 1 |

<2. Taxonomic Level >_<project>_<sex>_<wing-side>_<image_id>.TIF

**Table 2:**
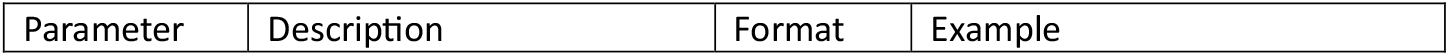
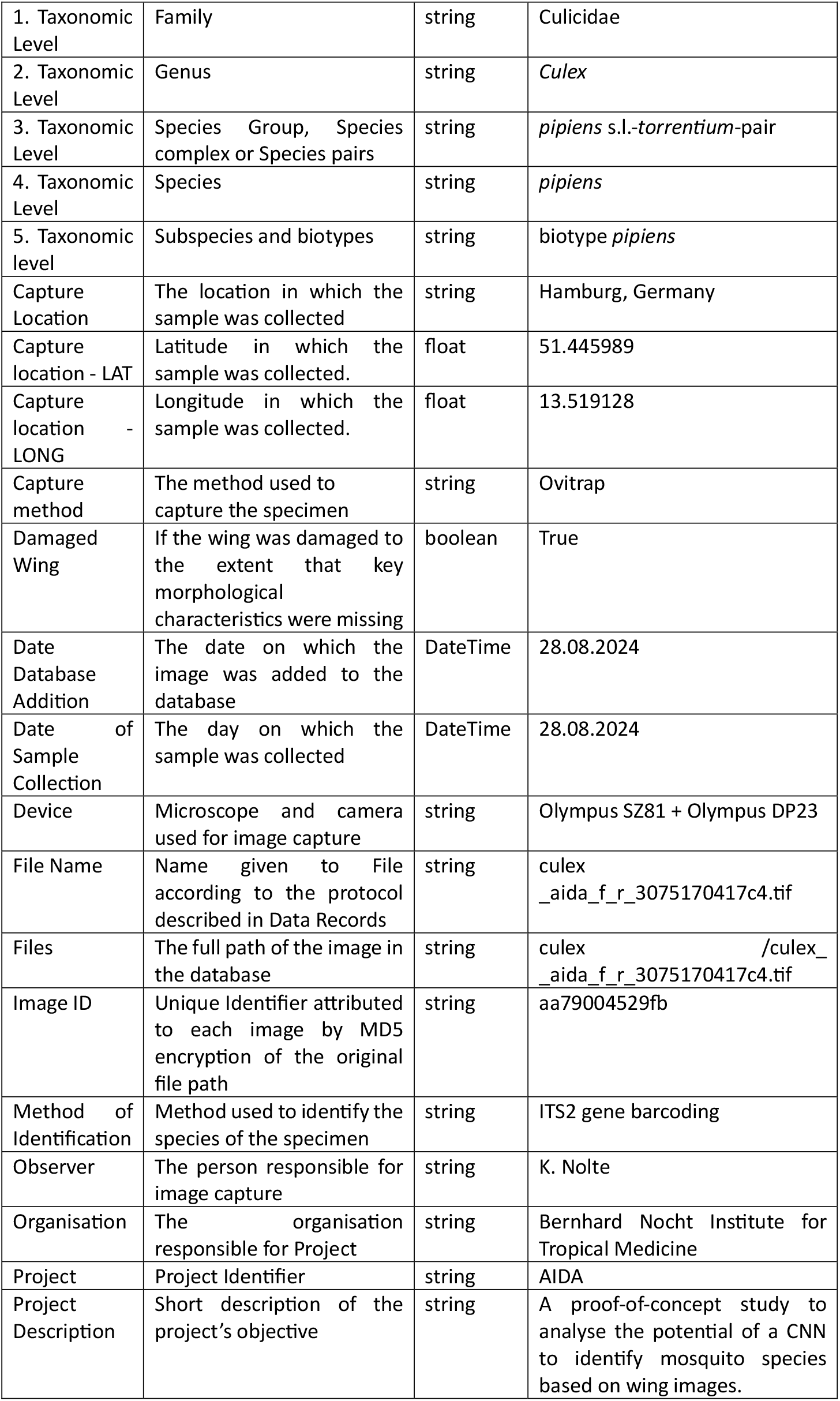
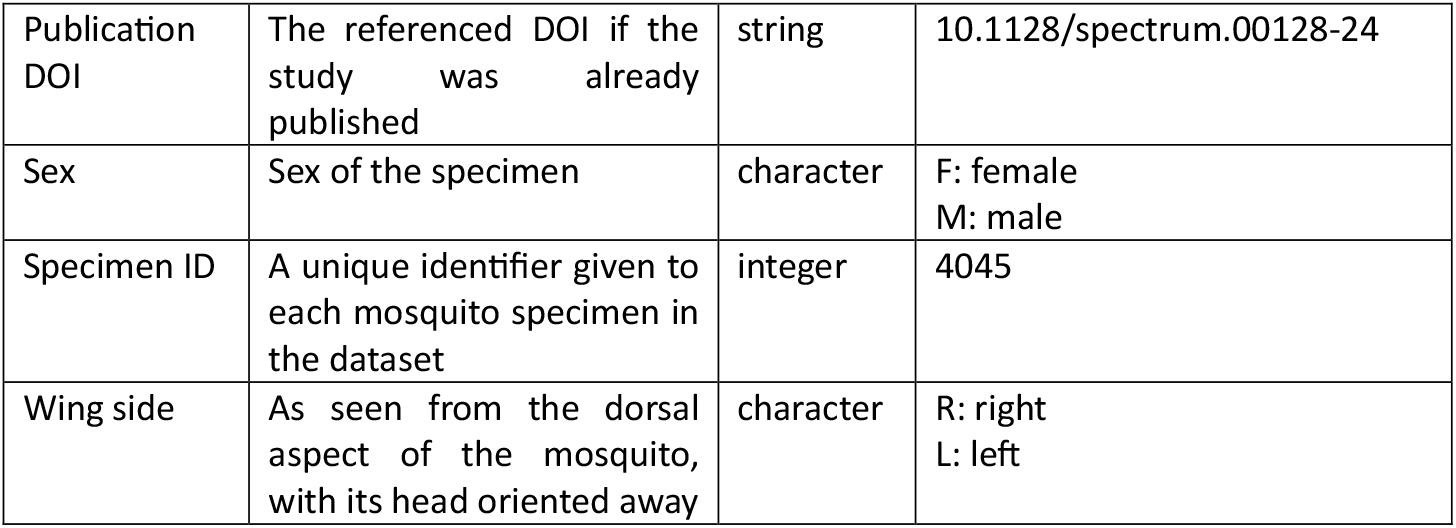
Description of meta parameters annotated to each image in the dataset.

### Technical Validation

Mosquito taxa were identified using mostly two approaches: by trained taxonomists with a dichotomous taxonomic key^4^ or through established molecular techniques such as gene barcoding or taxa-specific PCRs as described in the method section^24–26^.

The validity of the mosquito wing images has been demonstrated in previous studies. Specifically, several projects within this dataset, such as “landmark-mosquito-identification”^8^, “*aegypti*-diversity-study”^27^ and “landmark-*aedes*-collection”^9^ have shown that the wing images exhibit species-specific characteristics useful for wing geometric morphometrics for species identification or the analysis of population structure. Moreover, the data set has been proven valuable for deep learning applications, i.e. Sauer et al.^19^ (Project: CNN-study) and Nolte et al.^20^ (Project: ConVector) successfully trained CNN models to identify mosquito species. Moreover, Maciel-de-Freitas et al. (Project: *aegypti*-*Wolbachia*-study) demonstrated the utility of these image data to analyse the relationship between wing measurements, such as wing length and shape, and mosquito fitness parameters, such as fecundity, in *Ae. aegypti*^11^.

Despite extensive efforts to ensure comprehensive data collection, some metadata entries remain incomplete. Capture location is unavailable for 7.9% of samples, capture method is missing for 1.4%, and precise collection dates are missing for 24.1% of wing samples. Notably, the majority of samples lacking a collection date (75.5%) were derived from laboratory colonies.

Additionally, 48.4% of the wing images in this dataset originate from unpublished projects that adhered to data collection standards consistent with those in previously published studies.

### Usage Notes

This dataset supports the development and testing of geometric morphometric methods and machine learning models. However, users should be aware of certain limitations. The images were not captured under standardized conditions, resulting in significant variation across projects, such as differences in lighting and background (Fig. 2). This is due to the retrospective nature of data collection. Additionally, the dataset includes images of damaged wing samples (n=598), which lack certain morphological features.

**Figure 2.**
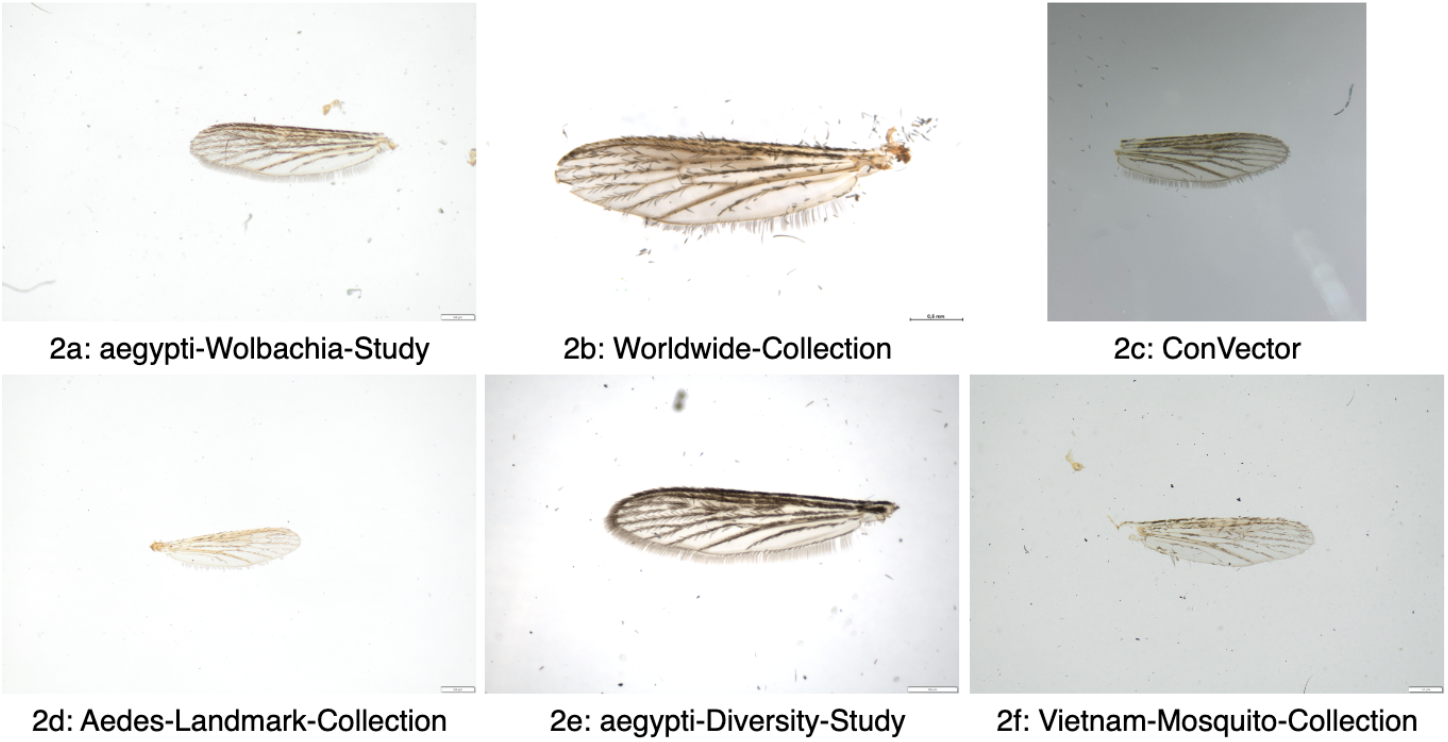
Example wing images of different projects of Ae. aegypti from the dataset. Images 2a, 2d, 2e and 2f were captured using the Olympus SZ61, 2b was captured using Leica M205c and 2c was captured using an iPhone SE with an attached micro-lens.

As further image data are collected for mosquitoes and other dipteran vectors, we plan to expand this dataset regularly. We also invite contributions from the scientific community to enhance this growing collection of wing images by contacting the authors.

## Data Availability

The dataset is available at BioImage Archive under CC BY 4.0 license https://doi.org/10.6019/S-BIAD1478

## Supporting information

Instructions to remove mosquito wings and capture images

## Acknowledgements

This project is funded through the Federal Ministry of Education and Research of Germany, with the grant number 01Kl2022, PAN-ASEAN Coalition for Epidemic and Outbreak Preparedness (PACE-UP; German Academic Exchange Service (DAAD) Project ID: 57592343), Federal Ministry of Health (ZMI1-2521NIK400), Deutsche Forschungsgemeinschaft (DFG) (grant number MA 9541/1-1 and SCHM 2413/9-1), Federal Office for Agriculture and Food (BLE) (FKZ 2819113519), Fundação Carlos Chagas Filho de Amparo à Pesquisa no Estado do Rio de Janeiro (grant number E-14/2019)), Heinrich-Böll-Stiftung grants for doctoral students of the German Federal Ministry of Education and Research, GIZ (Deutsche Gesellschaft für Internationale Zusammenarbeit; grant number 81281913), German Center for Infection Research, and Conservation, Building and Nuclear Safety (BMUB) through the Federal Environment Agency (UBA) (FKZ 3721484020).

## Author contributions

KN, FGS, RL wrote the manuscript. KN harmonized all data entries. All authors collected the original data. All authors revised and corrected the manuscript.

