## Supplementary material for "Comprehensive Mosquito Wing Image Repository for Advancing Research on Geometric Morphometric- and AI-Based Identification": Instructions to remove mosquito wings and capture images

### Wing Capture Guide for Dipteran vectors

#### Wing Removal

To remove the wings, you need two tweezers (or one dissecting needle and a tweezer) and a stereo microscope. One tweezer should have a very fine, pointed tip for grasping the wings, while the other is used to hold the insect during the process.

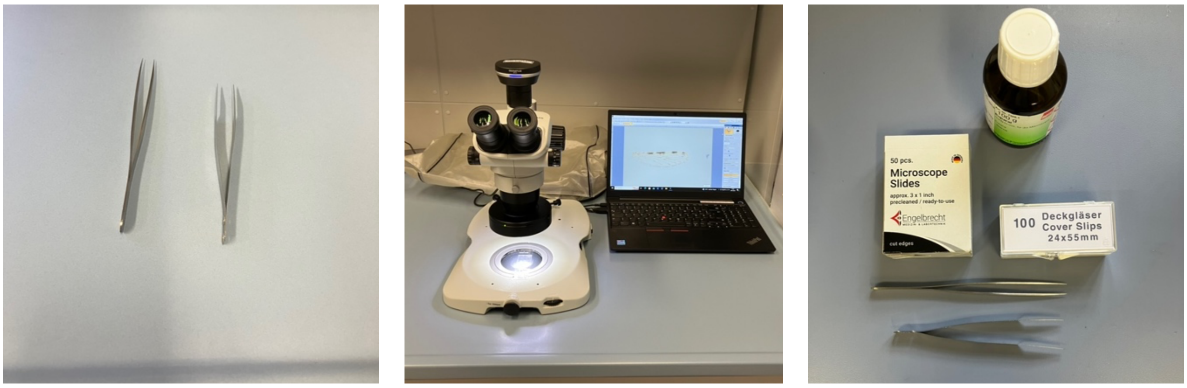


*Figure 1: Materials needed for wing removal. Two tweezers (left), microscope (middle) and optional materials needed for wing mounting: Euparal, microscope slides, cover slips and tweezers (right).*

Begin by using one tweezer (or the dissecting needle) to hold the insect body without applying pressure, ensuring stability without causing any deformation or destruction. Then target the junction between the thorax and the wing of the insect with the second tweezer. Pin the wing at this intersection with the pointed tweezer, taking care to avoid any contact with the wing. Once the wing is securely grasped, begin to release it from the body with gentle movements. It's crucial to handle the wing only at its junction from the body to minimize the risk of damage. Pull slowly on the wing until it is completely detached from the thorax.

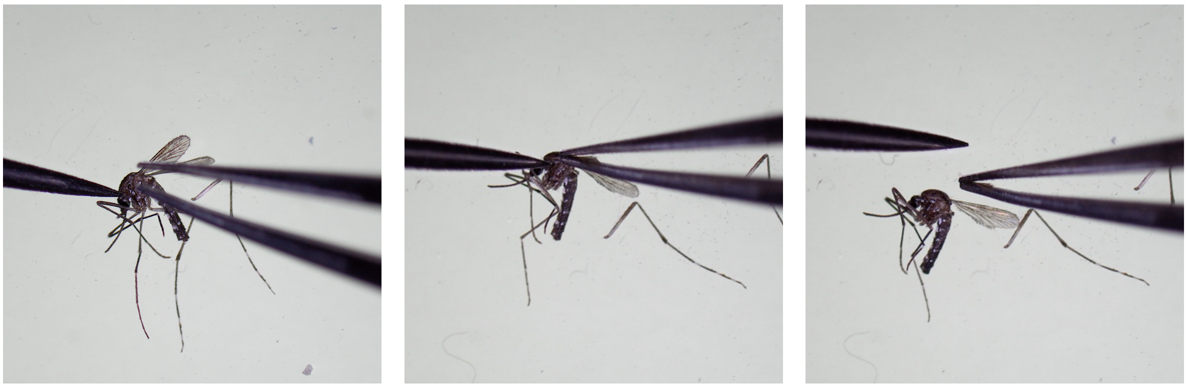

*Figure 2: Step by step wing removal. Fixating the mosquito (left), pinning the wing junction (middle) and wing removal (right).*

### **Wing Storage** (optional)

For long term storage, insect wings are mounted in an embedding medium such as Euparal (Carl Roth, Karlsruhe, Germany). Place a few small drops of the embedding medium onto a clean microscope slide as depicted in Fig. 3a. Then, carefully remove the insect wings and place them on the Euparal drop. Designate the left and right wings accordingly on the slide by positioning them on their respective sides, i.e. the left wing should be placed on the left side. The wing position in the drop should be corrected by the dissected needle. Press slightly the wing with the needle to the bottom of the drop, so it will be completely covered by liquid medium. To complete the process, gently position a cover slip (24 × 55 mm) over the wings, holding the cover slip from four corners by fingers. Try to drop the cover slip on wings from the very short distance from wings, simultaneously releasing the cover slip from four corners. This will prevent the formation of air bubbles.

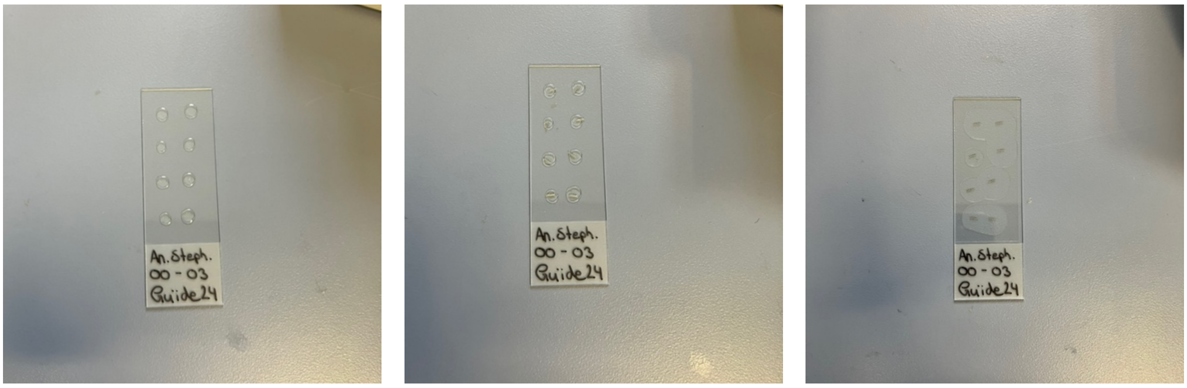


*Figure 3: Step by step preparation of insect wings for long term storage. Preparing microscope slide by placing small drops of Euparal (left), iteratively placing down the wings on the Euparal spots (middle) and putting on the cover slip (right).*

The slides have to be stored horizontally until the medium is dry. To speed up the drying process, it's recommended to place the slides in an oven set to a temperature between 50°C to 60°C for a minimum of one week, ideally extending to two weeks.

### Image Capture

When capturing images of the mounted wings, optimal illumination is essential for results. Ideally, utilize a stereo microscope equipped with a camera for image capture. If a microscope with a camera isn't available, a smartphone paired with a macro lens can serve as an alternative. Illuminate the wing from both the top and bottom to highlight the wing features effectively. You can achieve this by placing the slide on a petri dish. Place the wing on a neutral background. If the stereo microscope doesn't provide a neutral background, place a white sheet of paper beneath the slide. We use a 20x magnification when capturing images of insect wings. Be sure to record the magnification setting in the meta information (as described in the section below). Additionally, it's recommended to include a digital scale directly in the image whenever feasible, either by adding pixel size information to the meta data or incorporating a scale. When focussing on rather small insect species, a greater magnification can be considered. For the Olympus SZ61 (Olympus GmbH, Tokyo, Japan), the used magnification has to be set manually. Therefore, the scale should be calibrated by means of a microscope slide with a micrometer scale. This can be done measuring the micrometer scale using the “linear ruler” tool (Toolbar measure -> linear ruler).

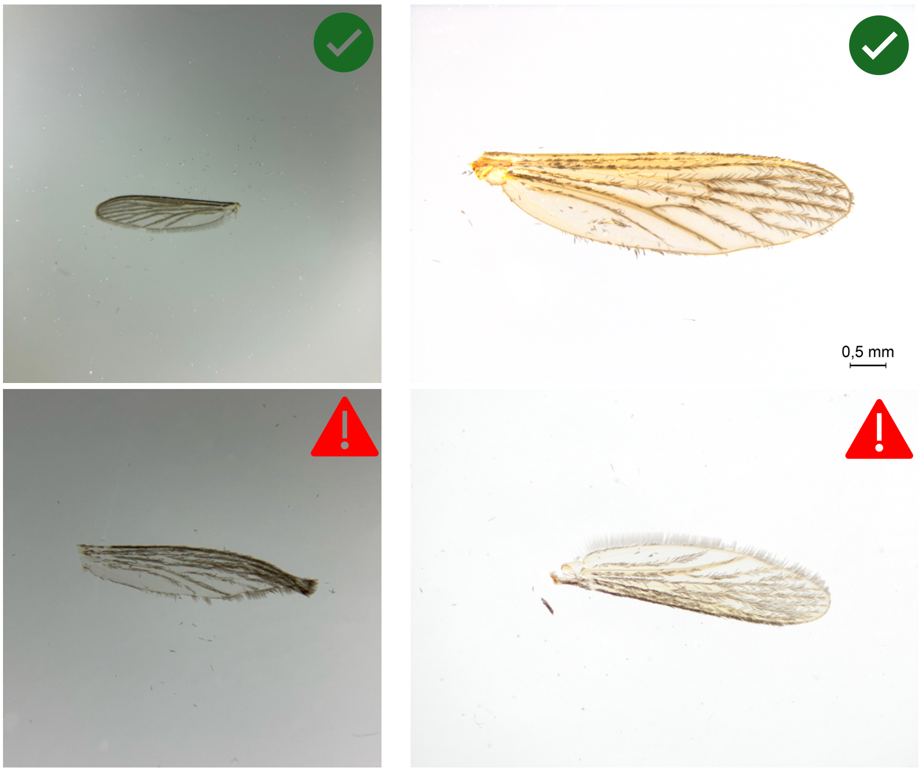


*Figure 4: Images captured using a smartphone with an attached macro lens (left), image captured with a camera attached to a stereo microscope, a scaled is burned in as size reference (right). Images suitable for classification are displayed at the top. Deformed, damaged or flipped wings are not suitable for classification (bottom). The latter can be corrected by rotating the image before the upload (bottom-right).*

### Data Recording

Files should be saved as either .PNG or .TIF files. For each data collection project, you should create a single folder, where all the images are stored, and which contains an excel file describing all the necessary meta information. Here is a excel template you can use.

| A | B | C | D | E | F | G | H | I | J | K | L |
|---|---|---|---|---|---|---|---|---|---|---|---|
| **File_Name** | **Specimen Nu** | **Genus** | **Species** | **Sex** | **Device** | **Wing Side** | **Date of Image Capture** | **Oberserver** | **Project** | **Location** | **Date of Sample Collection** |
| Autogenerated Name fo | If you use both | Genus of spe | Species of th | Sex of the sp | On what dev | Left or Right w | Date on which image w | Person who | As part of wh | Longitude / L | Date on which specimen wa |
| 1_aedes_albopictus_co | **1** | aedes | albopictus | f | Leice M320 | r | 20.09.23 | Kristopher N | convector | 9.99, 53.55 | 01.09.23 |
| 2_aedes_koreicus_conv | **2** | aedes | koreicus | f | Leice M320 | r | 07.09.23 | Kristopher N | convector | 9.99, 53.55 | 01.09.23 |
| 3_culex_pipiens_convec | **3** | culex | pipiens | f | Leice M320 | l | 21.09.23 | Kristopher N | convector | 9.99, 53.55 | 01.09.23 |
| 4_aedes_albopictus_co | **4** | aedes | albopictus | f | Leice M320 | r | 20.09.23 | Kristopher N | convector | 9.99, 53.55 | 01.09.23 |

Please name your files according to the template below. A name will also be generated by the Excel file for you (1st column, *File Name*) which you can copy. The file name in the excel template must align with image file name.

*<specimen-identifier>_<genus>_<species>_<project-name>_<wing side>*

| | | | |
|---|---|---|---|
| ConVector (Example) | Today at 15:25 | -- | Folder |
| 1_aedes_albopictus_convector_right.png | 20. Sep 2023 at 16:38 | 7,8 MB | PNG image |
| 1_aedes_koreicus_convector_left.png | 7. Sep 2023 at 15:03 | 10,2 MB | PNG image |
| 3_culex_pipiens_convector_right.png | 21. Sep 2023 at 09:31 | 2,7 MB | PNG image |
| 4_aedes_albopictus_convector_left.png | 20. Sep 2023 at 16:38 | 7,7 MB | PNG image |
| TemplateExcel (Example).xlsx | 20. Oct 2023 at 09:53 | 19 KB | Micros...k (.xlsx) |
